# TP53I11 Suppresses Extracellular Matrix-independent Survival and Mesenchymal Transition in Mammary Epithelial Cells

**DOI:** 10.1101/301499

**Authors:** Tongqian Xiao, Hai Zhang, Yuanshuai Zhou, Junsa Geng, Zhongjuan Xu, Yu Liang, Hong Qiao, Guangli Suo

**Affiliations:** CAS Key Laboratory of Nano-Bio Interface, Suzhou Institute of Nano-Tech and Nano-Bionics, Chinese Academy of Sciences, Jiangsu 215123, China; University of Chinese Academy of Sciences, Beijing 100049, China; University of Science and Technology of China, Anhui 230026, China; School of Life Sciences, Shanghai University, Shanghai 200444, China; Department of Molecular Biosciences, the University of Texas at Austin, Austin, Texas 78712, USA.

**Keywords:** TP53I11, metastasis, tumor progression, metabolism, AMPK

## Abstract

Extracellular matrix (ECM)-independent survival is an essential prerequisite for tumor metastasis and a hallmark of epithelial cancer stem cells and epithelial-mesenchymal transition (EMT). We found that, in MCF10A and MDA-MB-231 cells, loss of TP53I11 (Tumor Protein P53 Inducible Protein 11) enhanced the ECM-independent survival and suppressed glucose starvation induced cell death by increasing the activation of AMPK that confer cells metabolic flexibility to survive under stress conditions. We show here that, TP53I11 enhanced glycolysis and promoted proliferation of MCF10A and MDA-MB-231 cells in normal culture, but exerted negative effect on EMT, cell migration and invasion, and its overexpression suppressed tumor progression and metastasis of MDA-MB-231 cells *in vivo.* Considering cancer cells also are confronted with the hostile environment such as nutrient scarcity during tumorigenesis and metastasis, our findings suggested that the disruption of metabolic flexibility by TP53I11 through inhibiting AMPK activation resulted in the suppression of tumorigenesis and metastasis of breast cancer.

## INTRODUCTION

Epithelial cells is critically dependent on integrin-mediated attachment to extracellular matrix (ECM). The loss of detachment to ECM causes apoptotic cell death termed as “anoikis” (1). ECM deprivation-induced cell death involves intrinsic, extrinsic apoptosis (1), and autophagy (2). However, apoptosis inhibition has proven to be insufficient for maintaining ECM-independent survival in MCF10A mammary epithelial cells as a result of disruption in glucose utilization, ATP production and redox homeostasis (3–5), indicating that cellular metabolism plays a critical role in ECM-independent survival. Neoplastic transformation of epithelial cells is associated with the loss of anoikis and the gain of ECM-independent survival (6). The survival during ECM detachment has widely been considered a hallmark of epithelial cancer stem cells and oncogenic epithelial-mesenchymal transition (EMT) that is required for tumor metastasis (7–11).

For its pivotal role in carcinogenesis and metastasis, cellular metabolism has been regarded as a promising target for cancer therapy (12). A popular metabolic reprogramming described in tumor cells is the ‘Warburg effect’, in which cancer cells undergo increased aerobic glycolysis by an irreversible damage of mitochondrial function (13, 14). This theory is confronted with challenge from the emerging evidence suggesting that a critical subpopulation of cancer cells responsible for tumor maintenance, metastasis and stress resistance tend to rely more heavily on mitochondrial respiration than glycolytic catabolism for ATP generation (15–17). Furthermore, the ratio of oxidative phosphorylation (OXPHOS) to glycolysis was varied along with the tumor progress (18, 19). This flexibility of transition between glycolysis and OXPHOS enables cancer cells to better adapt to microenvironmental changes in different stages of tumor progression and metastasis (20).

TP53I11 (Tumor protein 53-inducible protein 11) was first reported two decades ago as one of early transcriptional targets of TP53 (21). So far, several studies about TP53I11 were reported mainly focusing on its function as a tumor suppressor to induce apoptosis in cancer cells (22, 23). But, the underlying mechanisms are still ambiguous. Our results show, for the first time, that although TP53I11 enhances epithelial cells aerobic glycolysis and proliferation in normal culture, it suppresses epithelial cell survival in ECM-detached and glucose starvation cultures, and reduces EMT, cell migration, tumor progression and metastasis of breast cancer cells. Our findings suggest that TP53I11 may function as a metabolic mediator of the oncogenic effect in breast cancer.

## RESULTS

### Loss of TP53I11 enhances ECM-independent survival by reducing the activation of AMPK

The expression level of TP53I11 in MCF10A is too low to be detectable in attached culture, but is markedly upregulated in detached culture (Fig. 1A). This indicates that TP53I11 may function on ECM-independent survival of mammary epithelial cells. To investigate this hypothesis, we first stably expressed the TP53I11 and TP53I11-shRNA in non-tumorigenic MCF10A cells (M10A-P11-OE and M10A-P11-KD, their vector control cells are M10A-V-OE and M10A-V-K, respectively) and in breast cancer MDA-MB-231 cells (M231-P11-OE and M231-P11-KD, their vector control cells are M231-V-OE and M231-V-KD, respectively) (Fig. 1B, C and Fig. 2A, B). Additionally, we generated homozygous TP53I11 knockout MCF10A cell line (M10A-P11-KO, its vector control cells is M10A-V-K) via CRISPR/Cas9 genome editing, and identified the mutation by sequencing (Fig. 1D). The knockdown and knockout of TP53I11 in MCF10A cells was validated via immunoblotting or qRT-PCR in detached culture (Fig. 1C and E). Furthermore, we found that loss of TP53I11 (Knockdown or knockout) promoted spheroid formation both in size (Fig. 1G and Fig. 2C) and number (Fig. 1H and Fig. 2D), increased the survival (Fig. 1I and Fig. 2E) and reduced anoikis (Fig. 1J, C, F and Fig. 2F) of detached MCF10A and MDA-MB-231 cells. Otherwise, loss of TP53I11 boosted, and gain of TP53I11 reduced the clonogenic survival of MCF10A cells subjected to 60 hours of detached culture (Fig. 1K); loss of TP53I11 stimulated no-lumen spheroid formation of MCF10A in Matrigel culture (Fig. 1L and M). For MDA-MB-231 cells, TP53I11 overexpression reduced the formation of branching spheroids, and inhibited the invasive growth of cells in Matrigel (Fig. 2G and H). Increased AMPK activation was reported to be implicated in anoikis resistance (24). Our results showed that loss of TP53I11 increased AMPK activation in detached MCF10A cells (Fig. 1N) and MDA-MB-231 cells (Fig. 2I). This indicates that loss of TP53I11 contributes to ECM-independent survival by promoting AMPK activation in cells.

**Fig. 1.**
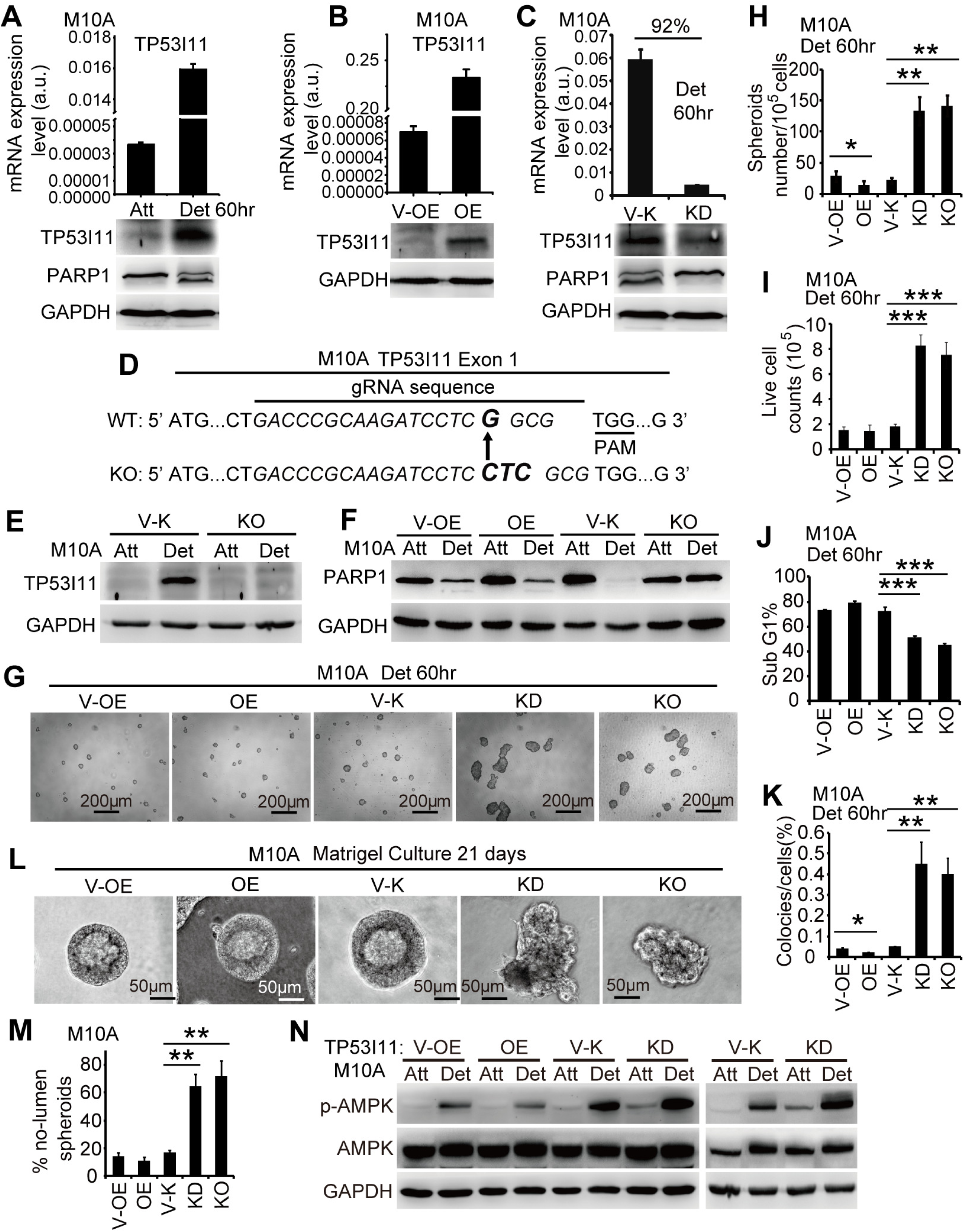
Loss of TP53I11 enhances ECM-independent survival in non-tumorigenic MCF10A mammary epithelial cells. (A) Upregulation of TP53I11 in MCF10A induced by 60 hours of detached culture, which is determined by qRT-PCR and immunoblotting assays. (B) TP53I11 was stably overexpressed, and (C) endogenous TP53IN was stably knocked down in MCF10A cells. The expression levels of TP53I11 were assessed by qRT-PCR and immunoblotting assays. (D) The sequence of TP53I11 mutation locus in monoclonal TP53I11 knockout MCF10A cell line. One base (G) was replaced by three bases (CTC) in exon 1 of TP53I11 genomic DNA. The gRNA sequence is shown in italics and the changed bases are shown in bold italics. The PAM sequence was underlined. (E) Immunoblotting assay to assess the knockout of TP53I11. (C) (F) PARP1 cleavage examined by immunoblotting assay to indicate apoptosis levels for MCF10A derived cells after 60 hours of detached culture. GAPDH was processed in parallel as an internal control for protein loading. (G) Representative images of aggregated spheroids of indicated MCF10A cells cultured in suspension for 60 hours. Scale bar = 200 μm. The number of spheroid bigger than 50 μm in diameter was quantified and the values were shown in (H). For indicated MCF10A cells after 60 hours detached culture, live cells remaining were counted by trypan blue exclusion and shown in (I); percent of SubG1 and survival colonies were shown in (J) and (K), respectively. (L) Representative bright field of images of Matigel culture of the indicated MCF10A cells. Scale bar = 50 μm. (M) The percentage of no-lumen spheroids of MCF10A indicated cells. (N) The AMPK phosphorylation levels of MCF10A indicated cells after 60 hours of detached culture were examined by Immunoblotting. GAPDH was processed in parallel as an internal control for protein loading. “Att”: attached culture; “Det”: detached culture; “a.u.”: arbitrary unit; “V-OE”: vector control of TP53I11 overexpression; “OE”: overexpression of TP53I11; “V-K”: vector control of TP53I11 knockdown or knockout; “KD”: TP53I11 knockdown; “KO”: knockout; “M10A”: MCF10A cells; “*”: *P* < 0.05, “**”: *P* < 0.01, “***”: *P* < 0.005, *P* values were determined using a two-tailed student’s *t* test; error bars represent S.D. from average value of at least three biological replicates.

**Fig. 2.**
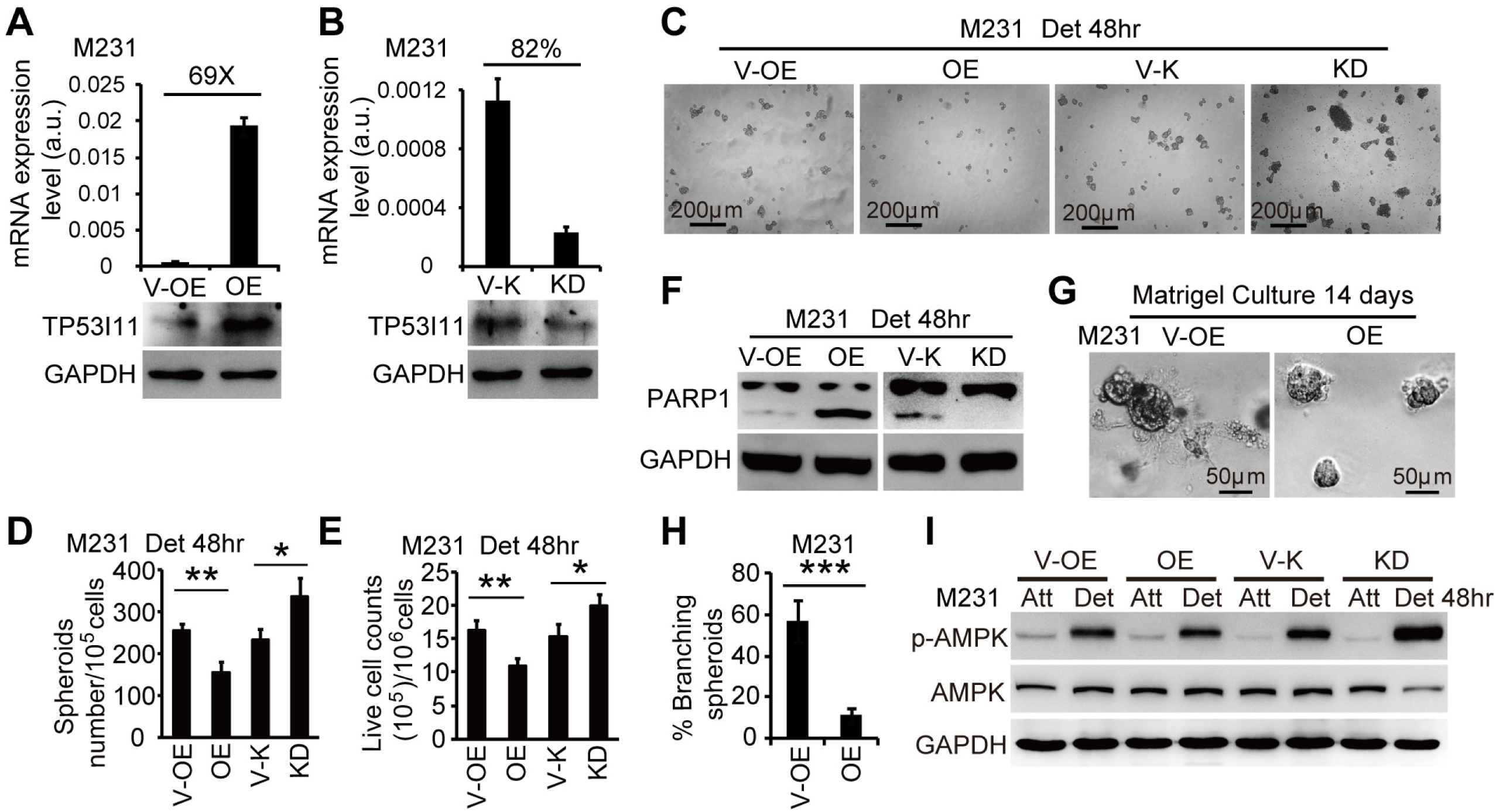
Loss of TP53I11 enhances ECM-independent survival in breast cancer MDA-MB-231 cells. (A) TP53I11 was stably overexpressed, and (B) Endogenous TP53III was stably knocked down in MDA-MB-231 cells. The expression levels of TP53I11 were assessed by qRT-PCR and immunoblotting assays. (C) Representative images of aggregated spheroids of MDA-MB-231 indicated cells cultured in suspension for 48 hours. Scale bar = 200 μm. The number of spheroid bigger than 50 μm in diameter was quantified and the values were shown in (D). (E) Live cells remaining after 48 hours of detached culture were counted by trypan blue exclusion. (F) PARP1 cleavage examined by immunoblotting assay to indicate apoptosis levels for MDA-MB-231 derived cells after 48 hours of detached culture. GAPDH was processed in parallel as an internal control for protein loading. (G) Representative bright field of images of Matigel culture of the indicated MDA-MB-231 cells. Scale bar = 50 μm. (H) The percentage of branching spheroids of MDA-MB-231 indicated cells. (I) The AMPK phosphorylation levels of MDA-MB-231 indicated cells after 48 hours of detached culture were examined by Immunoblotting. GAPDH was processed in parallel as an internal control for protein loading. “Att”: attached culture; “Det”: detached culture; “a.u.”: arbitrary unit; “V-OE”: vector control of TP53I11 overexpression; “OE”: overexpression of TP53I11; “V-K”: vector control of TP53I11 knockdown; “KD”: TP53I11 knockdown; “MDA231”: MDA-MB-231 cells; “*”: *P* < 0.05, “**”: *P* < 0.01, “***”: *P* < 0.005, *P* values were determined using a two-tailed student’s *t* test; error bars represent S.D. from average value of at least three biological replicates.

### TP53I11 promotes proliferation in normal culture, but reduces cell survival under glucose starvation by reducing the activation of AMPK

ECM-detached cells are subject to energetic stresses as a result of defects in glucose uptake (5, 24). For this reason, we examined the effect of TP53I11 on cell survival under glucose starvation. In normal cultures, the increase of PCNA expression level and cell viability indicates the positive impacts of TP53I11 on proliferation of MCF10A and MDA-MB-231 cells (Fig. 3A, C, D, K, M and N). Loss of TP53I11 decreased the cell number in G2/M phase, but increased that in G1/G0 phase (Fig. 3E and O). Opposite result was observed in TP53I11 overexpression cells (Fig. 3F and P). This indicates that loss of TP53I11 more likely induces the cells into quiescent stage (G0 phase) considering the corresponding decrease of proliferation. Under glucose starvation stress, we found that loss of TP53I11 decreased the PCNA expression, and improved cell survival of MCF10A (Fig. 3B, G and H) and MDA-MB-231 cells (Fig. 3L and Q).

**Fig. 3.**
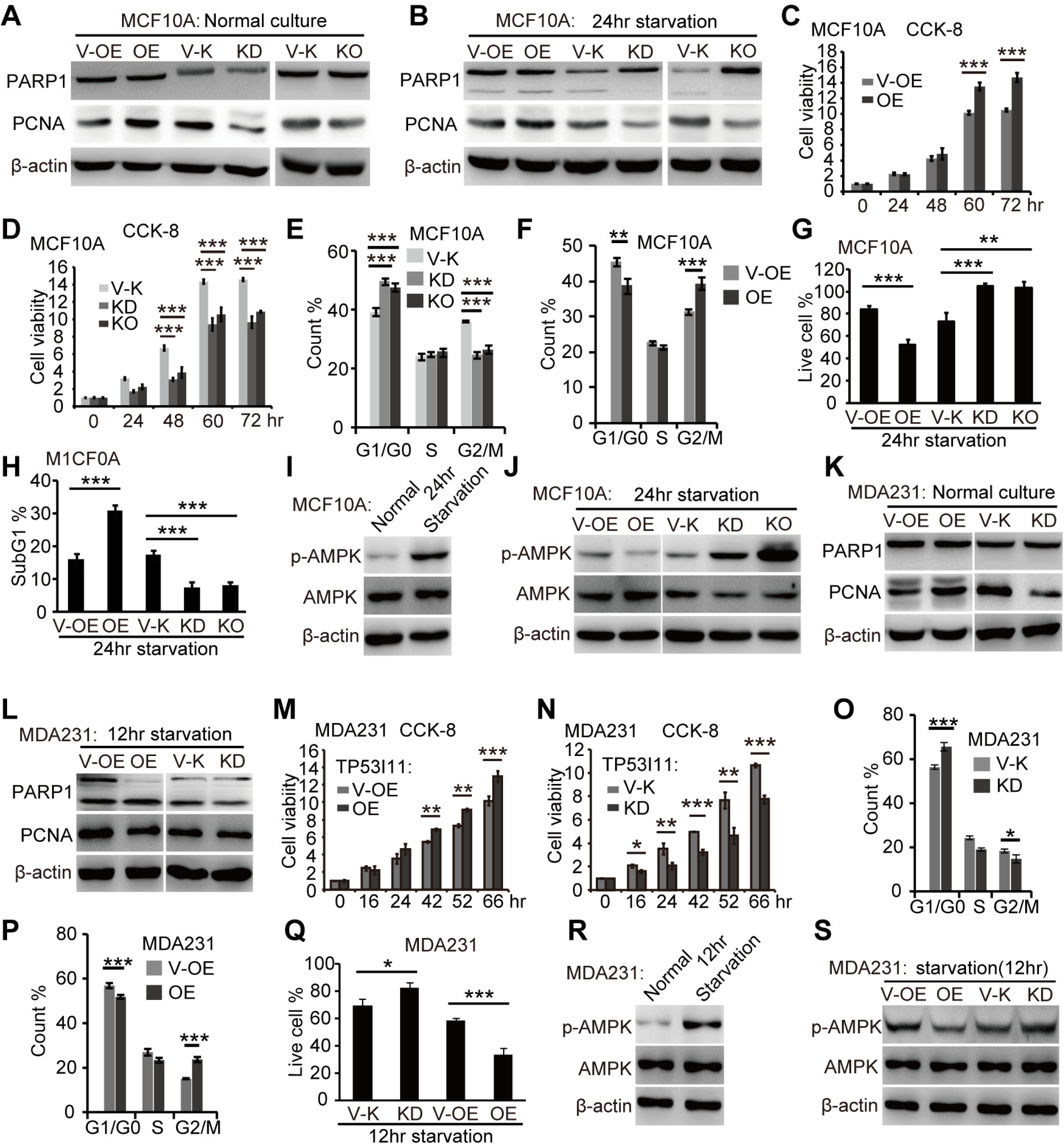
TP53I11 promotes proliferation in normal culture, but induces cell death under glucose starvation. Immunoblotting assay shows the expression of PCNA and PARP1 cleavage in indicated MCF10A cells (A) in normal cultures and (B) in glucose starvation cultures. (C and D) The viability of indicated MCF10A cells as a function of time. (E and F) Cell cycle distribution of indicated MCF10A cells. (G) The proportion of live cell and (H) SubG1 cell after 24 hours of glucose starvation. (I) Immunoblotting shows that the AMPK phosphorylation level is upregulated in MCF10A cells subjected to 24 hours of glucose starvation, and (J) the effects of TP53I11 on AMPK phosphorylation in indicated MCF10A cells subjected to 24 hours of glucose starvation. (K) Immunoblotting shows the expression of PCNA and PARP1 cleavage in indicated MDA-MB-231 cells in normal cultures and (L) in glucose starvation cultures. (M and N) The viability of indicated MDA-MB-231 cells as a function of time. (O and P) Cell cycle distribution of indicated MDA-MB-231 cells. (Q) The proportion of live cell after 12 hours of glucose starvation. (R) Immunoblotting shows that the AMPK phosphorylation level is upregulated in MDA-MB-231 cells and (S) the effects of TP53I11 on AMPK phosphorylation in MDA-MB-231 derived cells subjected to 12 hours of glucose starvation. “V-OE”: vector control for overexpression; “OE”: overexpression; “V-K”: vector control for knockdown or knockout; “KD”: knockdown; “KO”: knockout; “MDA231”: MDA-MB-231 cells; “*”: *P* < 0.05, “**”: *P* < 0.01, “***”: *P* < 0.005. *P* values were determined using a two-tailed student’s *t* test; error bars represent S.D. from average value of three biological replicates.

It was reported that AMPK could be activated to respond to metabolic stresses through preserving energy homeostasis in cells (25). We found that glucose starvation promoted AMPK activation in MCF10A and MDA-MB-231 cells (Fig. 3I and R). Under glucose starvation condition, overexpression of TP53I11 reduced, and loss of TP53I11 enhanced AMPK activation in both MCF10A and MDA-MB-231 cells (Fig. 3J and S). This indicates that TP53I11 affects the resistance of glucose starvation possibly via regulating AMPK activation in cells. Together, results in Fig. 3 suggest that loss of TP53I11 endows cell survival ability under glucose starvation through suppressing proliferation and improving AMPK activation.

### TP53I11 regulate the glycolysis and oxidative phosphorylation

It is well known that glycolysis plays a fundamental role in supporting cell proliferation (26, 27). To assess the effect of TP53I11 on glycolysis, we investigated extracellular acidification rate (ECAR) profiles of cells using the Seahorse instrumentation (Fig. 4A, B, I, M and Q). In MCF10A cells, loss of TP53I11 (knockdown or knockout) significantly decreased the levels of basal glycolysis, glycolytic capacity, and slightly decreased glycolytic reserve (Fig. 4C, D and E); in contrast, overexpression of TP53I11 significantly increased the levels of basal glycolysis, glycolytic and glycolytic reserve (Fig. 4F, G and H). The OCR results demonstrated that loss of TP53I11 decreased the levels of basal OCR (Fig. 5A, B), but increased the levels of maximal respiration and spare capacity (Fig. 5C and D). Overexpression of TP53I11 exerted no significant effect on basal OCR, maximal respiration and spare capacity (Fig. 5E, F, G and H). These results indicate that TP53I11 enhances glycolysis to contribute to cell proliferation. When glycolysis was inhibited by loss of TP53I11, cells rely on oxidative phosphorylation (OXPHOS) for energy compensation.

**Fig. 4.**
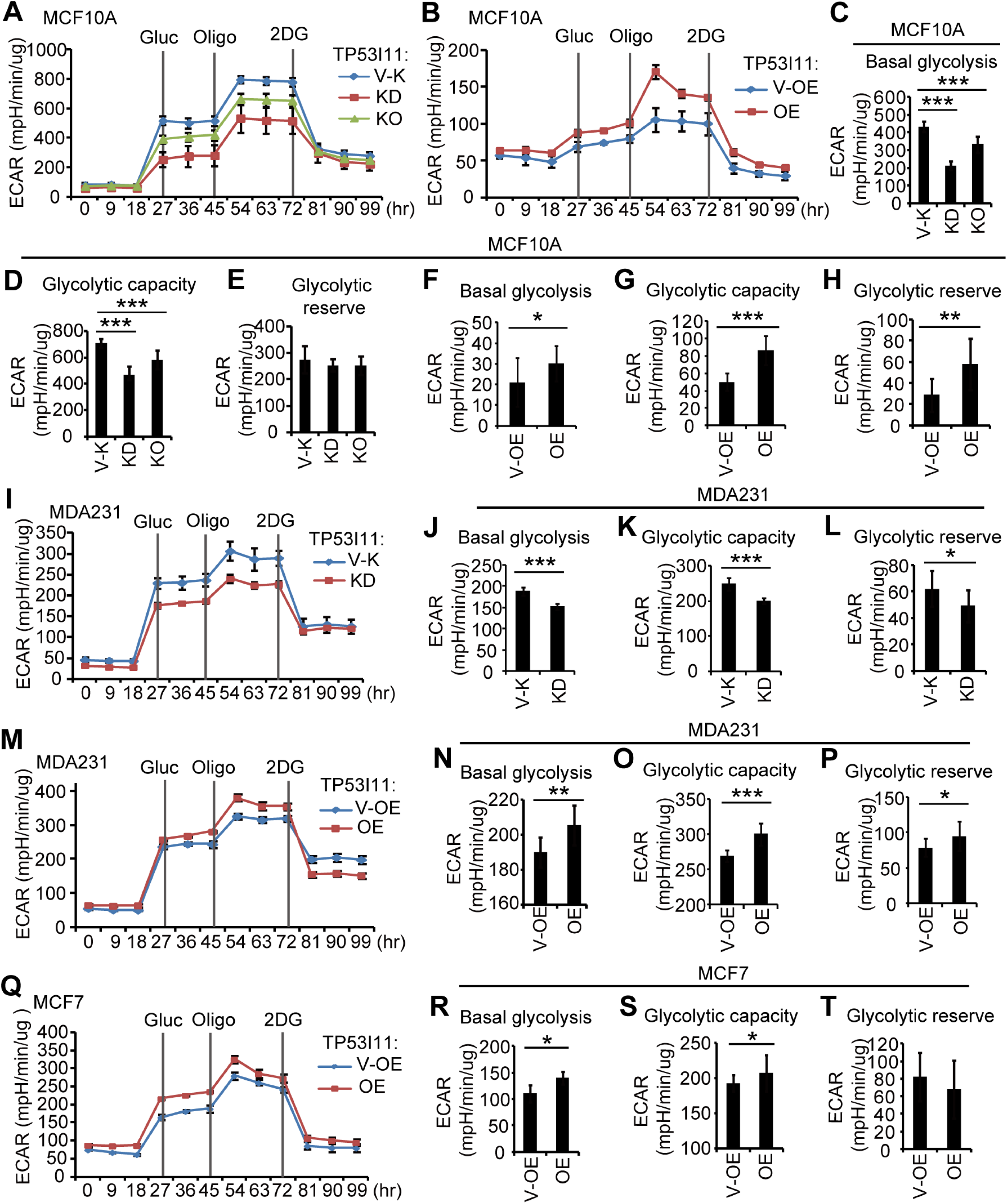
TP53I11 increases glycolysis in MCF10A, MDA-MB-231 and MCF7 cells. (A) Extracellular acidification rate (ECAR) of 10A-P11-KD(O) (KD or KO) and 10A-V-K (V-K) cells; n = 3; accordingly, the basal glycolysis, glycolysis capacity and glycolytic reserve are shown in (C) (D) and (E). (B) ECAR of 10A-P11-OE (OE) and 10A-V-OE (V-OE) cells; n = 3; the basal glycolysis, glycolysis capacity and glycolytic reserve are shown in (F) (G) and (H). (I) ECAR of 231-P11-KD (KD) and 231-V-K (V-K) cells; n = 3; the basal glycolysis, glycolysis capacity and glycolytic reserve are shown in (J) (K) and (L). (M) ECAR of 231-P11-OE (OE) and 231-V-OE (V-OE) cells; n = 3; the basal glycolysis, glycolysis capacity and glycolytic reserve are shown in (N) (O) and (P). (Q) ECAR of MCF7-P11-OE (OE) and MCF7-V-OE (V-OE) cells; n = 3; the basal glycolysis, glycolysis capacity and glycolytic reserve are shown in (R) (S) and (T). “V-OE”: vector control for overexpression; “OE”: overexpression; “V-K”: vector control for knockdown or knockout; “KD”: knockdown; “KO”: knockout; “Gluc”: glucose; “Oligo”: oligomycin; “Att”: attached culture; “Det”: detached culture; “MDA231”: MDA-MB-231 cells; “*”: *P* < 0.05, “**”: *P* < 0.01, “***”: *P* < 0.005. *P* values were determined using a two-tailed student’s *t* test; error bars represent S.D. from average value of three biological replicates.

**Fig. 5.**
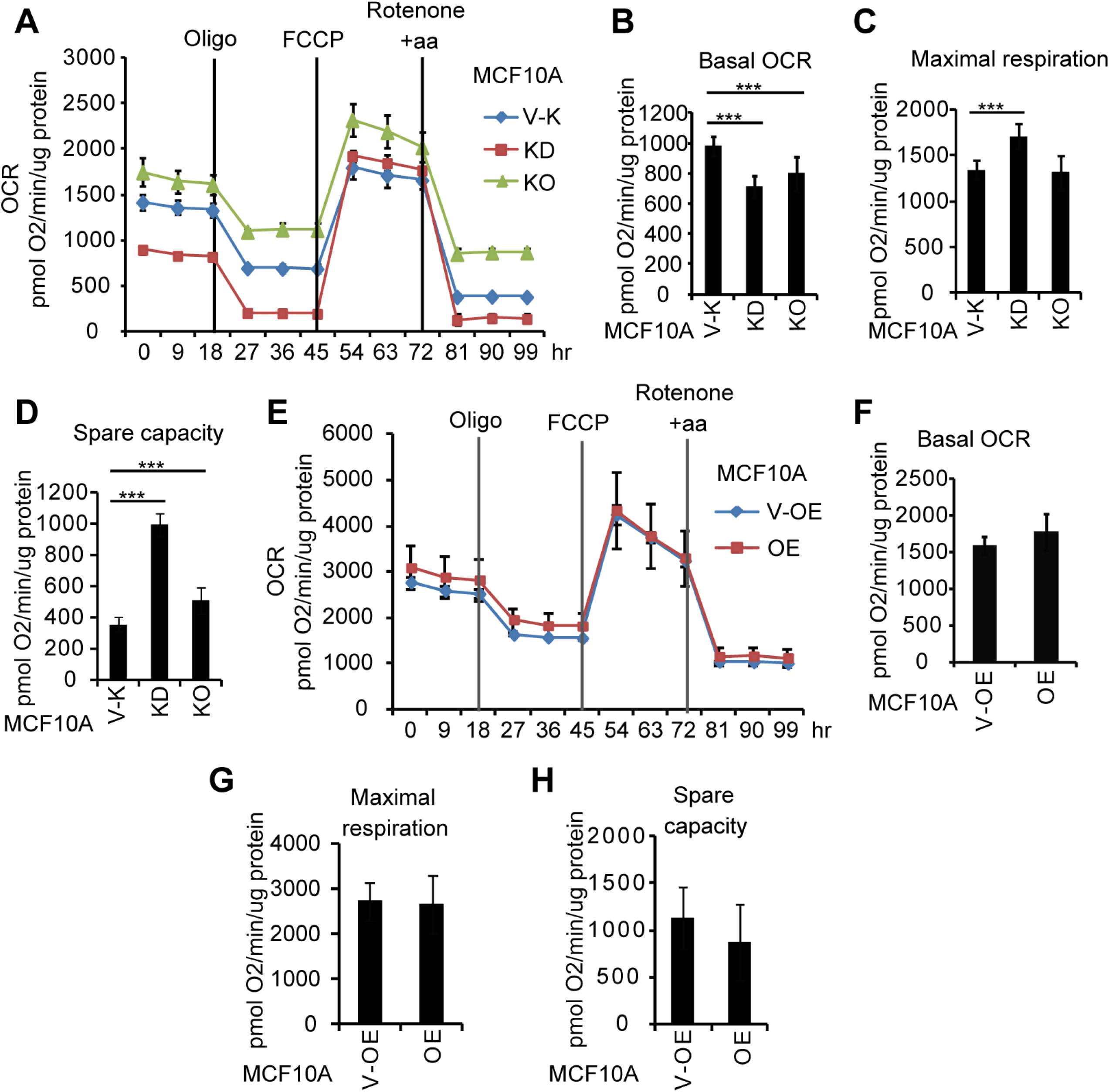
Loss of TP53I11 enhances Oxygen consumption rate (OCR) in MCF10A cells. (A) OCR of 10A-P11-KD(O) (KD or KO) and 10A-V-K (V-K) cells; n = 3; their basal OCR, maximal respiration and spare capacity are shown in (B), (C), and (D) respectively. (E) OCR of 10A-P11-OE (OE) and 10A-V-OE (V-OE) cells; n = 3; accordingly, their basal OCR, maximal respiration and spare capacity are shown in (F), (G) and (H), respectively. “V-OE”: vector control for overexpression; “OE”: overexpression; “V-K”: vector control for knockdown or knockout; “KD”: knockdown; “KO”: knockout; “Oligo”: oligomycin; “Rotenone + aa”: Rotenone + antimycin A; “***”: *P* < 0.005. *P* values were determined using a two-tailed student’s *t* test; error bars represent S.D. from average value of three biological replicates.

Similar ECAR results were found in MDA-MB-231 and MCF7 cells. Knockdown of TP53I11 significantly decreased, and overexpression of TP53I11 significantly increased the levels of basal glycolysis, glycolytic capacity and glycolytic reserve in MDA-MB-231 cells (Fig. 4J, K, L, N, O and P). TP53I11 overexpression significantly enhanced basal glycolysis and glycolytic capacity (Fig. 4R and S) in MCF7 cells, although there was no obvious effect on glycolytic reserve (Fig. 4T).

### Loss of TP53I11 promotes mesenchymal trail and cell migration

We noticed that M10A-P11-KD (knockdown) and M10A-P11-KO (knockout) cells displayed spindle morphology with increased cell scattering relative to M10A-V-K control cells in attached culture (Fig. 6A). To determine if these morphological changes were associated with EMT, we examined the effect of TP53I11 on several EMT markers in MCF10A and MDA-MB-231 (Fig. 6B and C). In MCF10A cells, loss of TP53I11 (knockdown and knockout) correlated with increased levels of mesenchymal markers, e.g., CDH2 (N-cadherin), VIM (Vimentin), SNAI2 (Slug) and SNAI1 (Snail) and decreased levels of epithelial markers such as CDH1 (E-cadherin), CLDN1 (claudin-1) and TJP1 (zona occludens 1, ZO-1) (Fig. 6B). In MDA-MB-231 breast cancer cells, TP53I11 overexpression reduced the expression of CDH2, VIM, SNAI2 and SNAI1, and enhanced the expression of CDH1, CLDN1 and TJP1 (Fig. 6C). These results showed a negative effect of TP53I11 on mesenchymal transition.

**Fig. 6.**
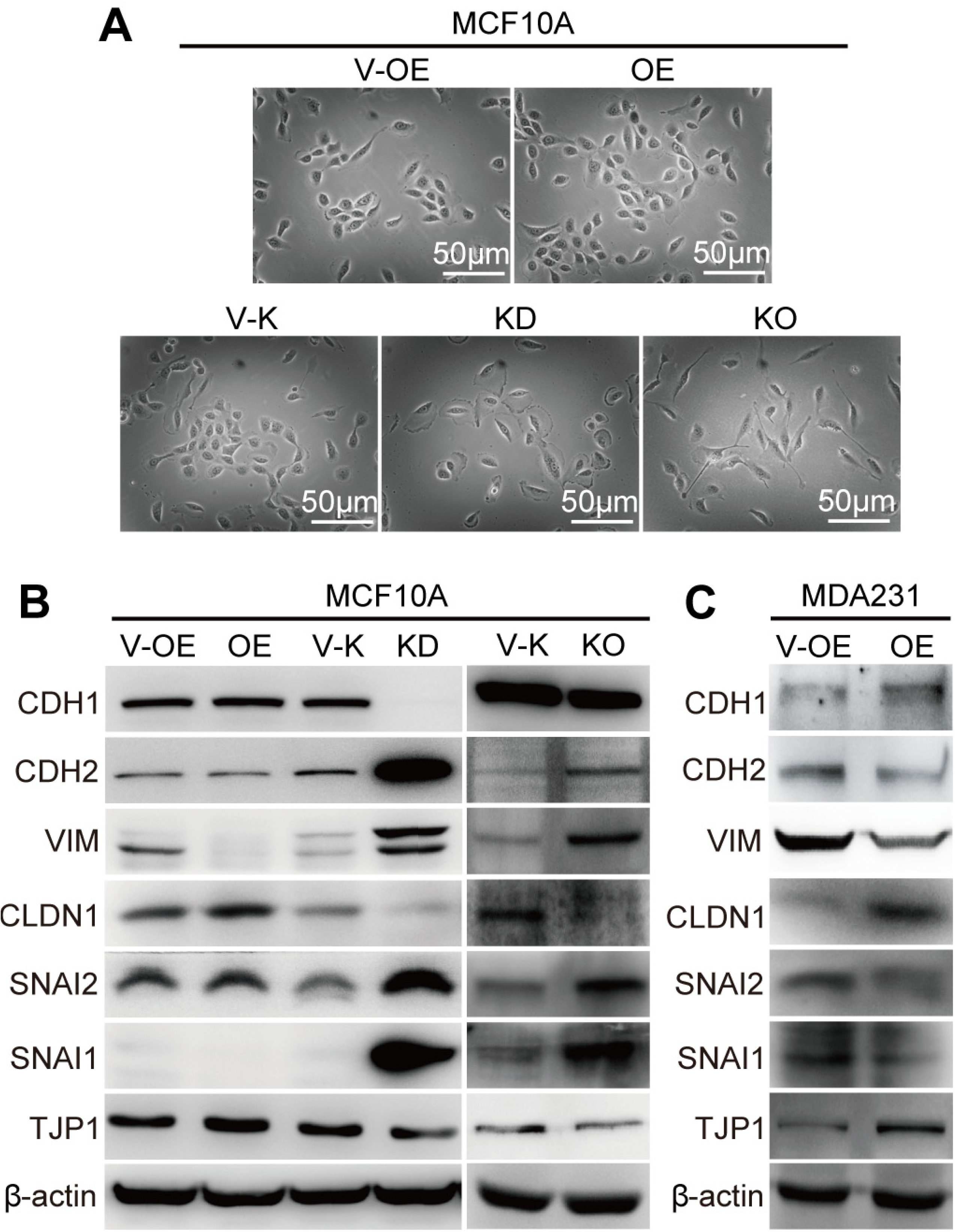
Loss of TP53I11 promotes mesenchymal traits. (A) Morphology of indicated MCF10A cells. Scale bar = 50 μm. Immunoblotting assay of EMT markers in (B) MCF10A and (C) MDA-MB-231 derived cell. “V-OE”: vector control for overexpression; “OE”: overexpression; “V-K”: vector control of knockdown or knockout; “KD”: knockdown; “KO”: knockout; “MDA231”: MDA-MB-231 cell line.

Since the mesenchymal transition is required for cell migration, we investigated the role of TP53I11 on migration and invasion in MCF10A and MDA-MB-231 cells. Loss of TP53I11 promoted and overexpression of TP53I11 suppressed wound closure, cell migration and invasion in MCF10A cells (Fig. 7A, B, C, D, E and F), and in MDA-MB-231 cells (Fig. 7G, H, I, J, K, L and M).

**Fig. 7.**
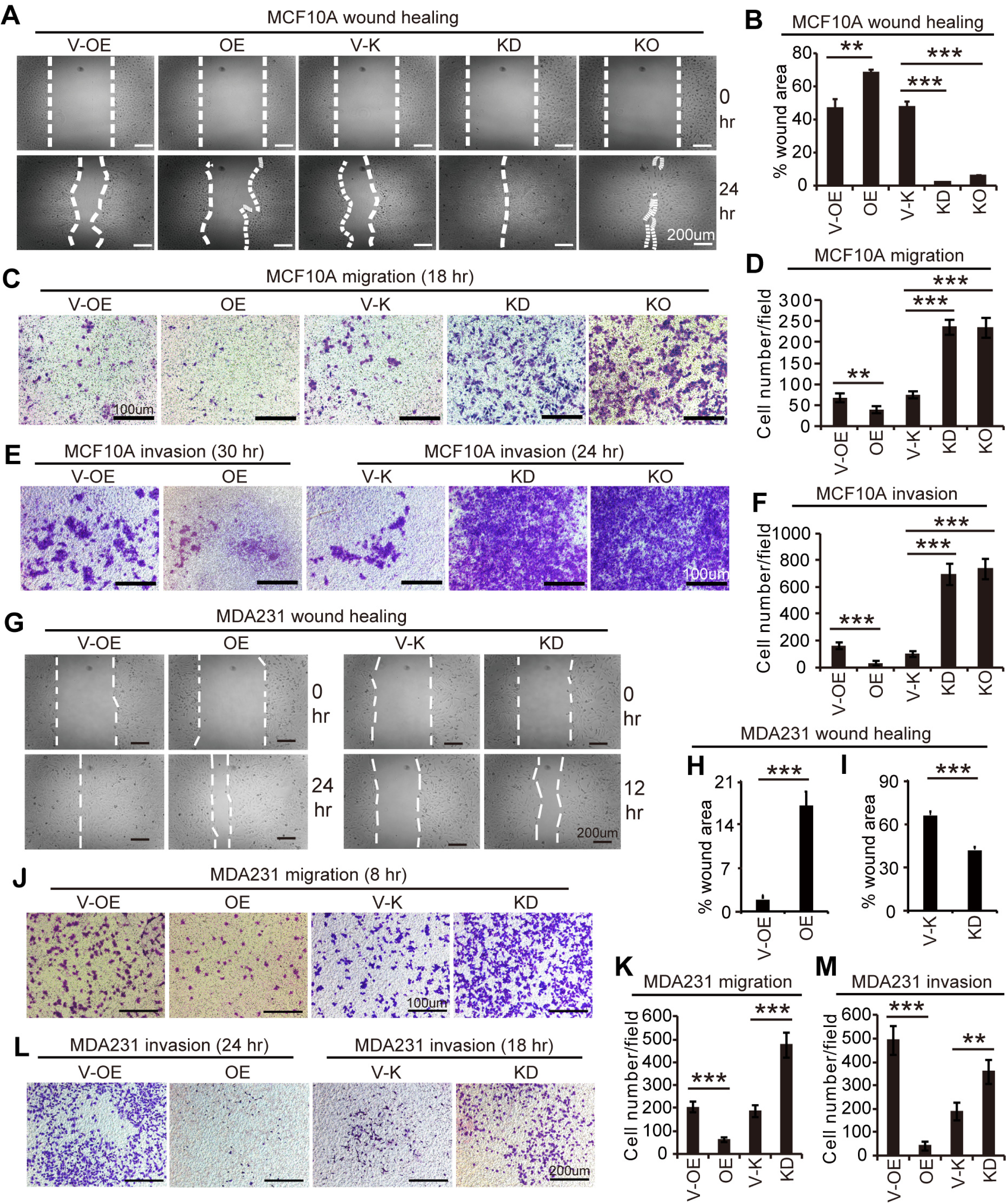
Loss of TP53I11 promotes cell migration and invasion. (A) Representative images of wound healing assays of MCF10A derived cells. The images were captured at indicated time of wounding. Dashed line represents the wound edge. Scale bar = 200 μm. (B) Percentage of wound healing at indicated time points. (C) Representative images of transwell migration assays, and (E) invasion assays of MCF10A derived cells. The images were captured at indicated time of cell culture. Scale bar = 100 μm. The number of migrated cells and invaded cells in each visual field were counted and show in (D) and (F), respectively. (G) Representative images of wound healing assays of MDA-MB-231 (MDA231) derived cells. The images were captured at indicated time of wounding. Dashed line represents the wound edge. Scale bar = 200 μm. (H and I) Percentage of wound healing at indicated time points. (J) Representative images of transwell migration assays, and (L) invasion assays of MDA231 derived cells. The images were captured at indicated time of cell culture. Scale bar = 100 μm in (J) and bar = 200 μm in (L). The number of migrated cells and invaded cells in each visual field were counted and show in (K) and (M), respectively. “V-OE”: vector control for overexpression; “OE”: overexpression; “V-K”: vector control of knockdown or knockout; “KD”: knockdown; “KO”: knockout; “**”: *P* < 0.01, “***”: *P* < 0.005. *P* values were determined using a two-tailed student’s *t* test; error bars represent S.D. from average value of at least three biological replicates.

### TP53I11 inhibit the tumor formation and metastasis in vivo

To evaluate the effect of TP53I11 on the tumor growth and metastasis *in vivo*, we transplanted M231-P11-OE (OE) and M231-V-OE (V-OE) cells into female nude mice (BALB/c) by orthotopic injection into the mammary fat pads or by tail-vein injection. We found that TP53I11 overexpression significantly reduced tumor growth in mammary fat pads as quantified by tumor volume and weight, and the tumors were only occurred in six of ten mices (Fig. 8A and B). H&E staining revealed three distinct layers in primer tumors: a necrotic tumor center (N) surrounded by a healthy, viable rim (V) with a layer of dermis (D) (Fig. 8C and E). The M231-P11-OE tumors generally demonstrated a smaller viable tumor rim relative to the control M231-V-OE tumors (Fig. 8C and E). Immunohistochemistry (IHC) staining of tumor sections showed that TP53I11 overexpression reduced the expression of mesenchymal markers CDH2 and VIM (Fig. 8F), decreased Ki67 expression and increased of caspase-3 activation (Fig. 8F and G). We found that TP53I11 overexpression substantially suppressed local invasion (Fig. 8C and D) and significantly reduced the metastasic burden in lungs (Fig. 8H and I). Colonization of the lungs by tail-vein injected cells was also measured by histology and staining for human VIM (Fig. 8J). Again, the TP53I11 overexpression significantly reduced the number and the size of tumor colonies in the lungs of tail-vein injected mice (Fig. 8K). These *in vivo* results confirmed of the *in vitro* findings that TP53I11 suppresses ECM-independent survival, cell migration, invasion and EMT.

**Fig. 8.**
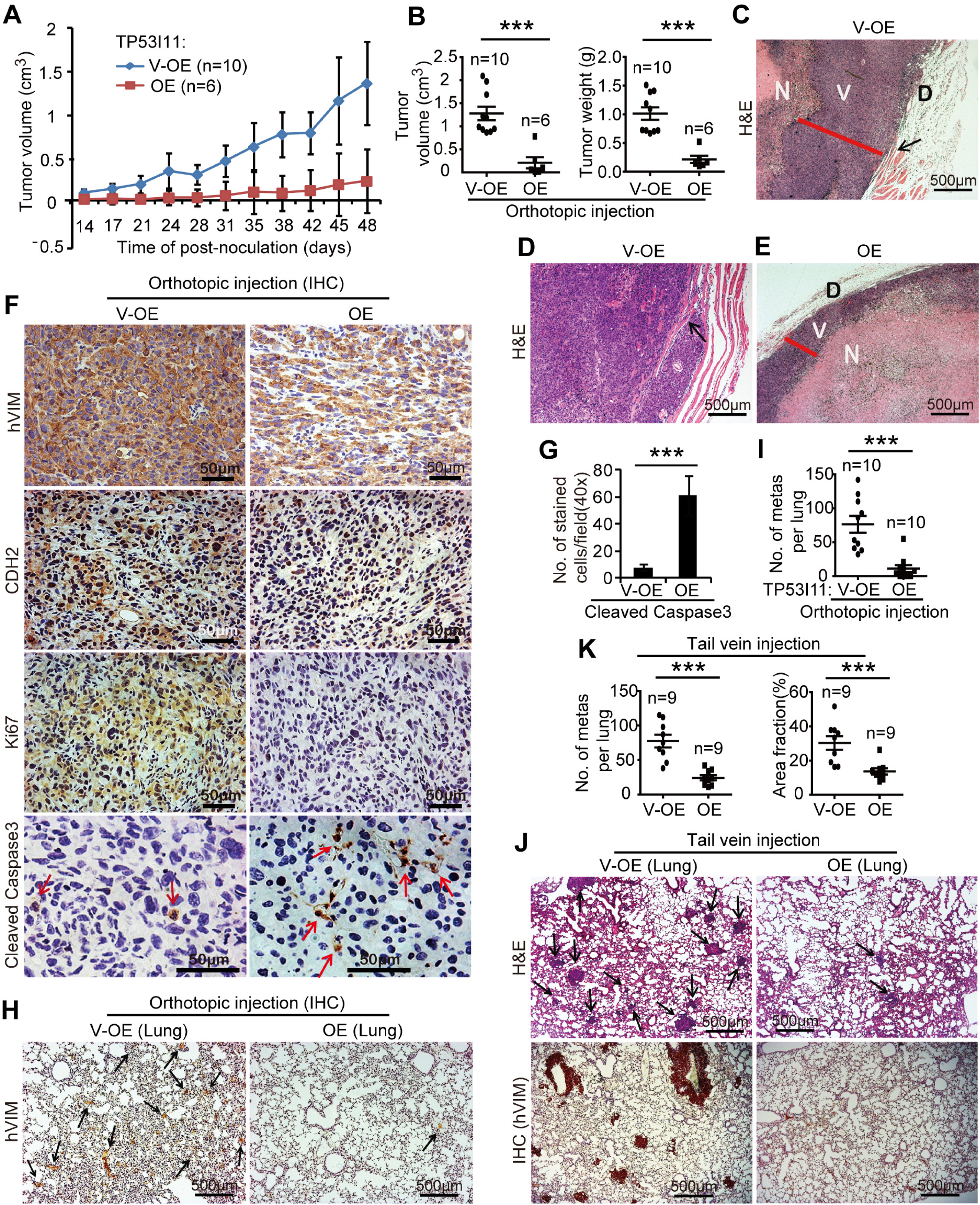
TP53I11 overexpression reduces MDA-MB-231 growth in mammary fat pads and metastasis to lung. Orthotopic injection: 231-P11-OE (OE) and vector control 231-V-OE (V-OE) cells were injected into the mammary fat pads of immunodeficient mice. (A) The tumor volumes were measured at the indicated days; (B) tumor volume and weight measured at day-48 in euthanized mice. (C, D and E) the tumors were sectioned and stained with haematoxilin and eosin (H&E); the necrotic center “N” is surrounded by a section of viable tumor rim “V” with an outer dermis layer “D”. TP53I11 overexpression reduced tumor invasion into nearly tissues. Black arrows point to the invasion locations. Red lines show the thickness of viable tumor rim. Scale bar = 500 μm. (F) Immunohistochemistry (IHC) staining for hVIM, CDH2, Ki67 and Cleaved Caspase3 in tumor sections; (G) number of Cleaved Caspase3-positive cells in each visual field; (H) IHC staining for hVIM in lung tissue sections and (I) hVIM-positive nodules in the lungs of mice from orthotopic injections. Tail vein injection: 231-P11-OE (OE) and vector control 231-V-OE (V-OE) cells were injected into immunodeficient mice through the tail veins and the mice were sacrificed after 35 days. (J) H&E and IHC (hVIM) staining of lung tissue sections; (K) the number and size of metastatic nodules. “V-OE”: vector control for overexpression; “OE”: overexpression; “***”: *P* < 0.005, *P* values were determined using a two-tailed student’s *t* test; results are expressed as mean ± SD from the indicated number of mice.

## DISCUSSION

This study was started from a finding that the expression of TP53I11 was markedly upregulated in MCF10A cells under ECM-detached culture. This motivated us to investigate if, and how TP53I11 functioned on ECM-independent survival, EMT and cell migration. For the first time, we reported that TP53I11 suppressed ECM-independent survival and EMT in MCF10A and MDA-MB-231 cells, and suppressed MDA-MB-231 cell growth in xenograft and metastasis to lung. This work supports other findings that TP53I11 was regarded as a tumor suppressor gene to promote cancer cells apoptosis induced by compounds (23, 28).

When MCF10A cells are subjected to detached culture, they lose viability with time most likely because of disruption in energy and redox metabolism. Neoplastic transformation of epithelial cells is invariably associated with the gain of ECM-independent survival. Ectopic expression of oncogenes, such as the ERBB2 receptor tyrosine kinase in MCF10A cells, has been shown to promote ECM-independent survival by stimulating ECM-independent glucose utilization (24, 29). Thereby, we investigated the effect of TP53I11 on glucose metabolism to clarify the mechanism of TP53I11 on ECM-independent survival. Our findings showed that overexpression of TP53I11 significantly increased the levels of glycolysis in MCF10A and MDA-MB-231 cells, and promoted their proliferation under normal culture condition. This is consistent with a popular theory that high level glycolysis allows the cells to balance their energy demands and supply the anabolic precursors for *de novo* nucleotide synthesis, and eventually increased the proliferation (27), which is the essence of “Warburg effect” to interpret tumor growth. However, we also found that TP53I11 suppressed cell survival in detached and glucose starvation culture, additionally, reduced the EMT and cell migration, and suppressed tumor expansion and metastasis of breast cancer cells *in vivo.* This suggests that TP53I11 plays a crucial role as a metastatic mediator in adapting cells to altered environments.

Recently, a growing body of evidence has been obtained suggesting that cancer-initiating cells/cancer stem cells are much more dependent on oxidative metabolism, which differ from the bulk tumor mass that prefer aerobic glycolysis (30, 31). It was reported that the ratio of OXPHOS to glycolysis was changed along with the tumor progress (18, 19). This metabolic flexibility endows cancer cells the survival advantage to cope with altered hostile environments during tumor progression and metastasis. The metabolic flexibility may be impacted or regulated by some crucial genes such as AMPK that plays a pivotal role in regulating metabolic flexibility to protect cells from hostile conditions (32–34). The loss or activation suppression of AMPK during tumorigenesis may sensitize tumor cells to apoptosis under hypoxic or nutrient depleted environments (32). Our results suggest that TP53I11 reduces AMPK activation and thereby disrupts the metabolic flexibility, which impairs cells survival ability under stressful growth conditions such as ECM-detached culture and glucose starvation. This disruption may be responsible for the suppression of tumor progression and metastasis during which cancer cells face with the hostile environment such as ECM detachment and nutrient scarcity. Given our results that support a tumor-suppressor function for TP53I11, it is possible that TP53I11 may function as a metabolic mediator of the oncogenic effect in breast cancer.

### Materials and Methods

#### Cell culture

MCF10A cell lines were obtained from ATCC (American Type Culture Collection, Manassas, VA, USA) and cultured in growth medium containing Dulbecco’s Modified Eagle Medium/F12 (Gibco) supplemented with 5% horse serum, 20 ng/ml EGF, 10 μg/ml insulin, 500 ng/ml hydrocortisone, 100 ng/ml cholera toxin, and 1% penicillin/streptomycin (35). MDA-MB-231 and MCF7 (ATCC, Manassas, VA, USA) cells were grown in complete Dulbecco’s Modified Eagle Medium (DMEM) (Gibco) supplemented with 10% fetal bovine serum, and 1% penicillin/streptomycin under humidified conditions in 95% air and 5% CO_2_ at 37°C.

#### Plasmids construction, viral production and infection

The cDNA of human TP53I11 was cloned from MCF10A cells by RT-PCR with primers (Table 1) and inserted into pQCXIH retrovector (Clontech Laboratories, Mountain View, CA). MCF10A and MDA-MB-231 cells were infected with TP53I11 retrovirus and control vector for 48 hr and selected with 100 μg/ml hygromycin B. The shRNA for TP53I11 knockdown (Table 1) was cloned into pLKO.1 vector to generate TP53I11-shRNA. Cells were infected with TP53I11-shRNA lentivirus for 48 hr and selected with 1 μg/ml puromycin. The sgRNA of TP53I11 (Table 1) was constructed into lenti-CRISPR plasmid (Addgene plasmid 49535) (36). The CRISPER-cas9-mediated TP53I11 knockout in MCF10A cells was constructed according to the method from previous report (36). Overexpression or knockdown of genes was evaluated by qRT-PCR with primers shown in Table 1 and immunoblotting.

**Table 1.**
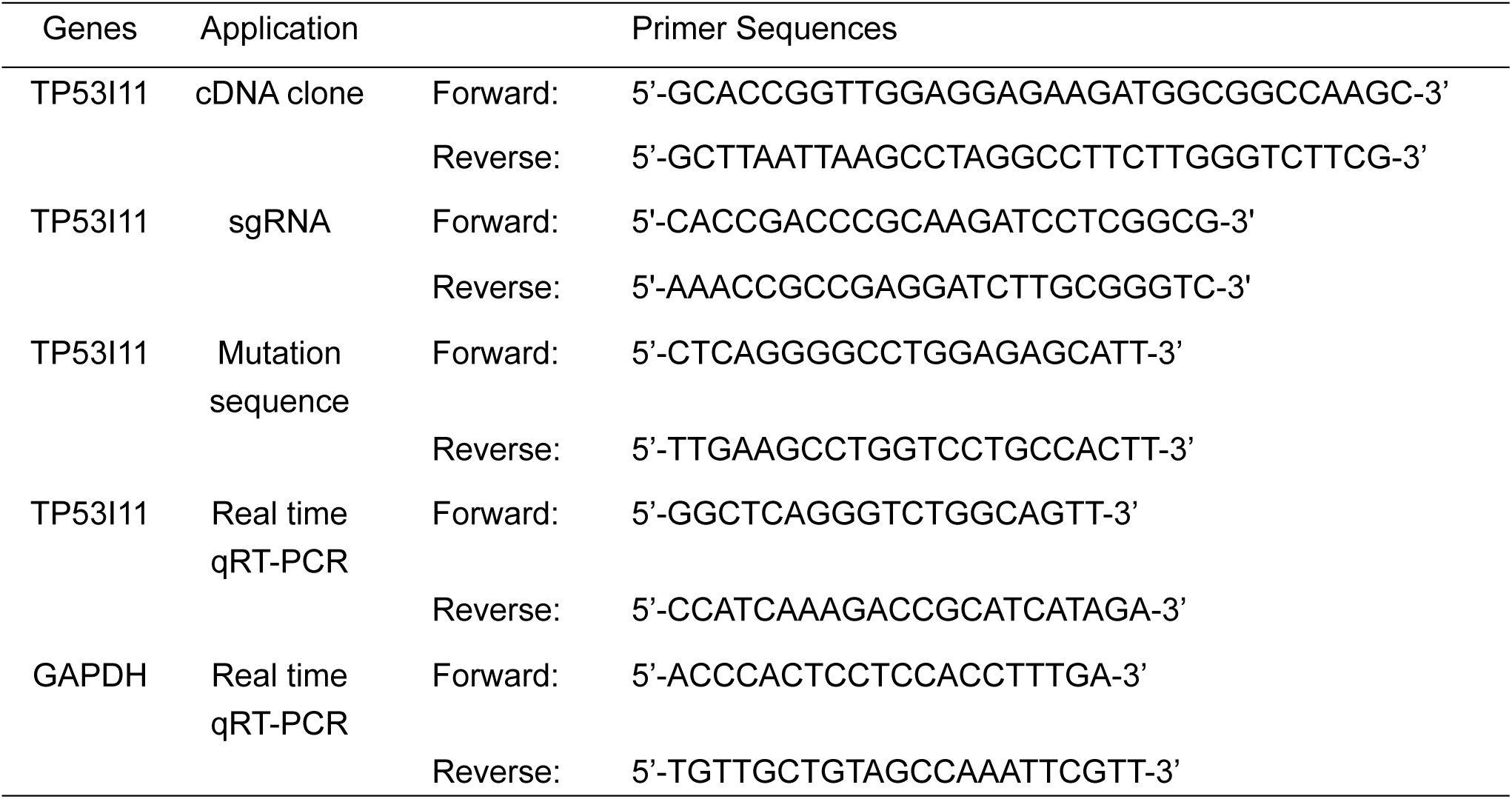
Primer sequences used in the study

#### ECM-detached culture and survival assay

Cells were performed ECM-detached culture in low detached dishes coated with Poly-HEMA to prevent cell adhesion (37). Briefly, trypsinized MCF10A and MDA-MB-231 cells were seeded at a concentration of 1 × 10^6^ cells per 100-mm low-detached dish and cultured in growth medium containing 0.5% methylcellulose to avoid survival effects caused by clumping of cells. The suspended cells were cultured at 37°C in a 5% CO_2_ atmosphere for 60 hours. Then, the number of aggregated spheroids with diameter more than 50 μm for MCF10A and MDA-MB-231 was counted. The cell viability was detected by typan blue exclusion test and clonogenic survival assay. Cell apoptosis was assessed with SubG1 assay and analysis of PARP1 by Immunoblotting.

#### Matrigel culture

MCF10A cells were cultured in growth factor reduced Matrigel using the proposal protocol from the Brugge laboratory (35). Briefly, 8-well chamber slides (Millipore) were chilled and coated with 45 μl/well of Matrigel (BD Biosciences). The Matrigel coat was solidified by incubation at 37°C for 30 minutes prior to cell seeding. Trypsinzed MCF10A cells were seeded at a density of 6,000 cells/well in 400 μl of medium containing 2% Matrigel. This medium was changed every other day, and the bright field images of the Matrigel cultures were acquired on days 21 post-seeding. The culture of MDA-MB-231 cells in Matrigel was performed using a modified protocol developed for MCF10A 3D culture. Cells were mixed in complete DMEM containing 2% Matrigel and then seeded on a coat of previously solidified Matrigel with 4,000 cells/well in 8-well chamber. The medium was changed every other day, and bright field images of the cultures were acquired by microscopy (Nikon eclipse TS100) on days 14 post-seeding.

#### Cell proliferation assay

Under normal attachment-culture condition, proliferation rate was calculated by the ration of live cell number at indicated culture time point over live cell number at zero culture time point. Cell Counting Kit-8 (CCK-8) (DOJINDO) allows sensitive colorimetric assays for the determination of cell viability in cell proliferation assay. Under normal attached culture, CCK-8 was used for cell proliferation assay according to the manufacturer’s instruction. For this assay, cells were seeded into a 96-well plate (6,000 cells/per well for MCF10A derived cell lines, 5,000 cells/per well for MDA-MB-231 derived cell lines) and cultured in growth medium or complete DMEM medium before assay. The optical density (OD) of each well at 450 nm was recorded on by VICTOR X4 Mutilabel Plate Reader. The proliferation rate was calculated by the ration of OD value at indicated culture time point over that at zero culture time point.

#### Immunoblotting

Cell lysates were prepared in lysis buffer (25 mM pH 7.4-Tris, 150 mM NaCl, 1 mM EDTA, 10% glycerol, 1% Nonidet P-40, 0.1% SDS, 1% sodium deoxycholate) supplemented with 1 x protease cocktail. The lysates were cleared by centrifugation at 21,000 x g for 15 minutes at 4°C, and proteins in the supernatants were quantitated using the Lowry Protein Assay quantitation Kit (Bio-Rad). Lysates (40 μg) were fractionated on 10% SDS-PAGE gels and transferred to polyvinylidene difluoride (PVDF) membranes. The membranes were incubated with primary antibodies followed by incubation with goat anti-rabbit or anti-mouse HRP-conjugated secondary antibody. Finally, the membranes were visualized by chemiluminescence following standard protocols and photographed by Fujifilm LAS 4000 luminescent image analyzer (Fujifilm Life Science).

#### RNA extraction and quantitative real-time polymerase chain reaction (qRT-PCR)

Total RNAs were extracted from cells using the RNAeasy Mini Kit (QIAGEN) and reverse transcribed using the ABI reverse transcription kit according to the manufacturer’s instructions. The reaction of real time PCR was performed in Power SYBR Green PCR Master Mix (Life Technologies Corporation, Carlsbad, CA, USA) on the ABI PRISM 7500 Sequence Detection System (Applied Biosystems). The relative expression levels were expressed in arbitrary units (a.u.) where the Ct value of the gene of interest was normalized to that of GAPDH. The primers for qRT-PCR were designed using the Primer Express software (version 2.0, Applied Biosystems) and showed in Table 1.

#### Glycolysis assay

Rates of glycolysis were determined by measuring extracellular acidification rate (ECAR) using a Seahorse XF24 Extracellular Flux Analyzer with the Seahorse XF Glycolysis Stress Test Kit (Seahorse Bioscience, Billerica, MA, USA) according to the manufacturer’s instruction. The day prior to the assay, cells were seeded in 24-well culture plate from Seahorse Bioscience at density 5 × 10^4^ cells/well. Prior to the assay, growth medium was replaced with serum -free XF Seahorse Assay Medium supplemented with 2 mM glutamine, and then cells were incubated for 1 hour at 37°C in an incubator without CO_2_. Cartridges equipped with oxygen- and pH-sensitive probes had been preincubated with calibration solution overnight at 37°C in an incubator without CO_2_. At the time of measurement, cells were then placed in the XF24 Extracellular Flux Analyzer with cartridges of probes. ECAR was evaluated in a time course before and after injection of the following compound: (i) glucose (10 mM final concentration); (ii) oligomycin (1 μM final concentration); (iii) 2-deoxyglucose (2-DG) (10 mM final concentration). The ECAR measurements were normalized to the total protein amount per well. All the measurements were replicated in three times.

#### Measurement of mitochondrial respiration

The day prior to the assay, MCF10A-derived cells were seeded in 24-well assay plate (Seahorse Bioscience, North Billerica, MA, USA) at a density 4×10^4^ cells/well and cultured in growth medium. Prior to the assay, medium was replaced with serum-free XF Seahorse Assay Medium (Seahorse Bioscience) supplemented with 1 mM sodium pyruvate and 25 mM glucose, and then cells were incubated for 1 hour at 37°C in an incubator without CO_2_. Cartridges equipped with oxygen- and pH-sensitive probes had been preincubated with calibration solution overnight at 37°C in an incubator without CO_2_. At the time of measurement, cells were then placed in the XF24 Extracellular Flux Analyzer with cartridges of probes. OCRs were evaluated in a time course before and after injection of the following compounds: oligomycin (1 μM final concentration), FCCP (Carbonyl cyanide-4-(trifluoromethoxy) phenylhydrazone (1 μM final concentration) and Rotenone + antimycin A (AA) mixure (10 μM each final concentration). The Oxygen consumption rate (OCR) measurements were normalized to the total protein amount per well. All the measurements were replicated three times.

#### Transwell migration and Matrigel invasion assays

MCF10A or MDA-MB-231 derived cells were starved for growth factors overnight, and then plated in the top upper chambers of the transwell (8 μm pore size; Corning) and cultured in growth medium. To each bottom lower chamber, 500 μl growth medium was added as attractant of migration. At the indicated time, cells on the basolateral microporous membrane were fixed with 100% methanol and stained with 0.1% crystal violet solution. Three independent migration experiments each with duplicate technical repeats per sample were performed. For the invasion assays, each upper chamber was first coated with 50 μl basement membrane growth factor-reduced Matrigel, then seeded with cells and incubated for indicated hours. The number of cells on the basolateral microporous membrane was determined as described above for the transwell migration assay.

#### Metastasis assays

All mice were housed in an exclusion barrier facility under protocol from the Rutgers Institutional Animal Care and Use Committee (IACUC). For orthotopic inoculation, cells (6 × 10^6^ in 60 μl) were injected orthotopically into the mammary fat pad of 6-week-old female nude mice. Tumors were measured every 4 days and tumor volume was calculated according to the formula [volume = πLW^2^/6] (38), where “L” is the length of xenograft and “W” is the width of xenograft. All animals were sacrificed at 48 days after the initial implantation. Primary tumors, lung tissues were collected and fixed in 4% paraformaldehyde for further analysis. For tail inoculation, 1 × 10^6^ viable cells were injected into the lateral tail vein. All animals were sacrificed at 35 days after inoculation. Lung tissues were collected, fixed in 4% paraformaldehyde for further analysis. Paraformaldehyde-fixed tumors and lungs were performed haematoxylin and eosin (H&E) staining and immunohistochemistry (IHC) assay using antibodies such as anti-N-cadherin (CDH2), anti-human vimentin (hVIM), anti-Ki67 and anti-cleaved caspase3.

#### Statistical analysis

Statistical significance was determined by unpaired two-tailed student’s *t*-test and one-way ANOVA. *P* < 0.05 was considered statistically significant. The average or mean data were calculated using Microsoft Excel or GraphPad Prism 5 software.

## ACKNOWLEDGEMENTS

We thank Jianwu Dai, Qiangbin Wang, Zhijun Zhang, Renjun Pei and Guosheng Cheng for insightful comments on the study, Xiang He for her excellent technical assistance. Nano-Bio-Chem Centre in Suzhou Institute of Nano-Tech and Nano-Bionics (SINANO) is acknowledged for professional assistance of cell imaging and FACS assay. This work was supported by National Natural Science Foundation of China (Grant No. 31471307); and Ministry of Science and Technology (MOST) (Grant No. 2017YFA0104301 and 2014CB965003). G. Suo was also supported by Hundred Talent Program of Chinese Academy of Sciences.

## CONFLICT OF INTEREST

The authors declare no conflict of interest.

